# Mapping malaria by combining parasite genomic and epidemiologic data

**DOI:** 10.1101/288506

**Authors:** Amy Wesolowski, Aimee R. Taylor, Hsiao-Han Chang, Robert Verity, Sofonias Tessema, Jeffrey Bailey, T. Alex Perkins, Daniel Neafsey, Bryan Greenhouse, Caroline O. Buckee

## Abstract

Recent global progress in scaling up malaria control interventions has revived the goal of complete elimination in many countries. Decreasing transmission intensity generally leads to increasingly patchy spatial patterns of malaria transmission, however, and control programs must accurately identify remaining foci in order to target interventions efficiently. In particular, mosquito control interventions like bed nets and insecticide spraying are best targeted to transmission hotspots, and the role of connectivity between different pockets of local transmission becomes increasingly important since humans are able to move parasites beyond the limits of mosquito dispersal and re-introduce parasites to previously malaria-free regions. Quantifying the connectivity between regions due to human travel, measuring malaria transmission intensity in different areas, and monitoring parasite spatial spread are therefore key issues for policy-makers because they underpin the feasibility of elimination and inform the path to its attainment. To this end, recent efforts have been made to develop new approaches to incorporating human mobility into spatial epidemiological models, for example using mobile phone data, and there has been a surge of interest in collecting spatially informative parasite samples to measure the genomic signatures of parasite connectivity. Due to their complicated life-cycles, *Plasmodium* parasites pose unique challenges to researchers in this respect and new methods that move beyond traditional phylogenetic and population genetic tools must be developed to harness genetic information effectively. Here, we discuss the spatial epidemiology of malaria in the context of transmission-reduction interventions, and the challenges and promising directions for the development of integrated mapping, modeling, and genomic approaches that leverage disparate data sets to measure both connectivity and transmission.

## Main Text

### The spatial dimensions of malaria control and elimination strategies

Assessing variation in the spatial and temporal distribution of infection, or in the distribution of a particular pathogen phenotype like drug resistance, is an important prerequisite for any infectious disease control effort. For malaria, these considerations are critical across the range of transmission settings (Figure 1). In pre-elimination settings for example (e.g. E-2020 countries including Swaziland, Costa Rica, China, and South Africa [1]) surveillance programs must locate and keep track of imported infections, conduct contact tracing, and ensure that onward transmission resulting from importation events are extinguished rapidly. For countries with intermediate transmission (e.g. Bangladesh, Namibia, Kenya), control programs must identify the transmission foci contributing to infections in the rest of the country and locate importation hotspots, since these will require different levels of transmission reduction versus surveillance and treatment efforts. Even in high transmission settings (e.g. Uganda, Nigeria, Democratic Republic of Congo, Myanmar), which have traditionally focused on monitoring clinical cases and scaling up control and treatment strategies across the country, the renewed interest in measuring transmission has also raised the possibility of more effective program evaluation, to assess the impact of interventions on transmission in different regions. Of particular importance, in moderate to high transmission settings, coordination between different regions is likely to be important when human mobility between them is frequent.

**Figure 1:**
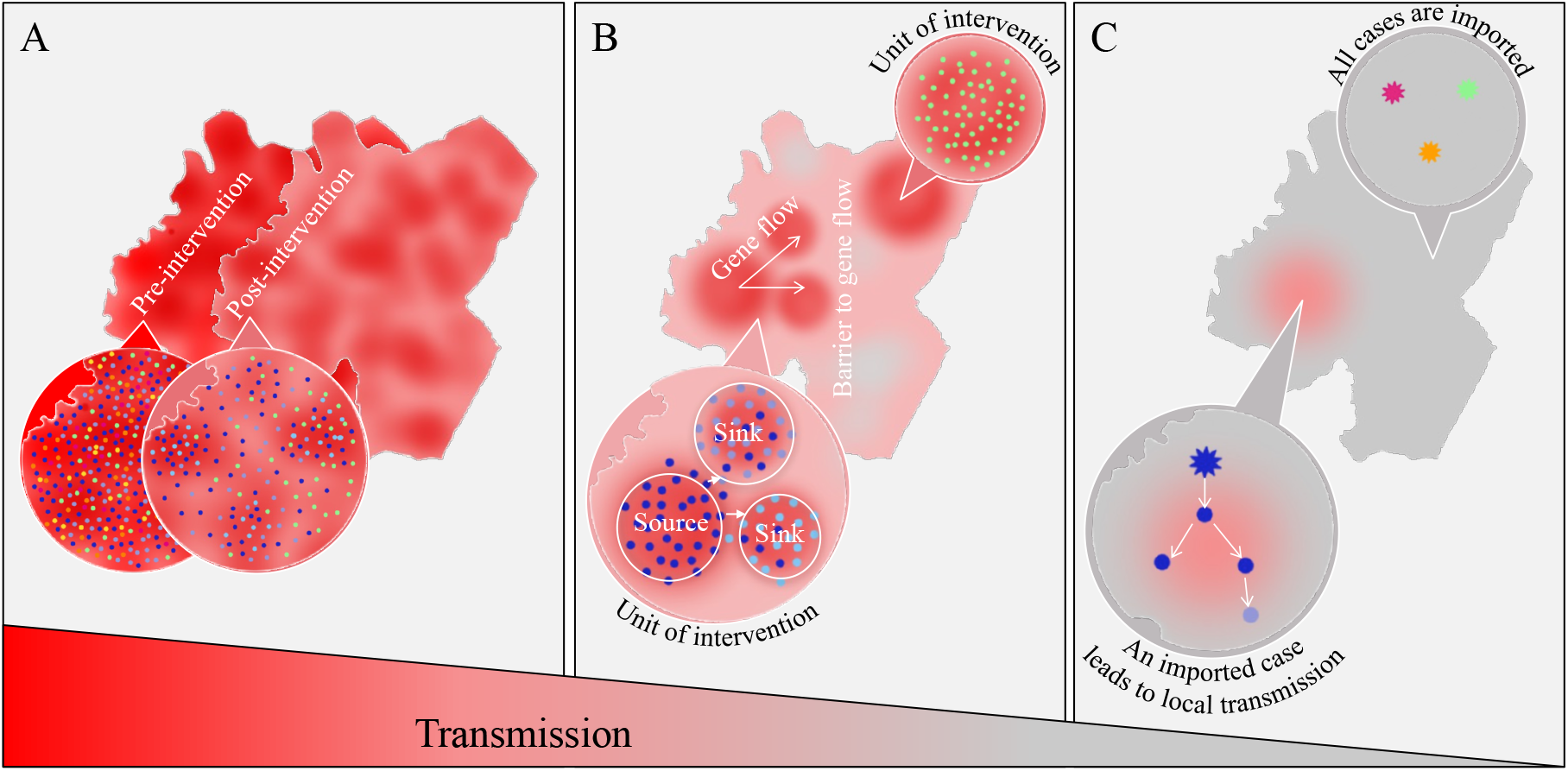
Actionable insight from genetic epidemiological studies of malaria across a range of transmission settings. This schematic depicts actionable insight that can be obtained from genetic epidemiological studies of malaria across a range of transmission settings, from high transmission (pink) on the left to low transmission (grey) on the right. Here both imported (stars) and local (points) infections are shown that may origin from different parasite lineages (various colors). In high transmission settings, parasites mix panmictically, polyclonal infections are common, and the goal is to evaluate the effectiveness of ongoing interventions Genetic correlates of declining transmission (e.g. diversity) can provide sensitive indicators of the impact of an intervention. At intermediate transmission, parasites may cluster into interconnected populations. The goal is to delineate regions into units for targeted intervention, and to identify sources that seed transmission for maximally efficient resource allocation. In this setting, models incorporating human mobility and genetic measures of parasite relatedness can provide directional estimates of connectivity between parasite populations. At very low transmission most infections are imported. The goal is to identify origins of imported parasites, quantify any onward transmission and, if onward transmission exists, the average length of local transmission chains. Models incorporating detailed case data, including genetic data and travel history, can reconstruct transmission chains to infer who acquires infection from who and how (WAIFW).

### Epidemiological models of malaria spatial epidemiology

A variety of modeling approaches have been used to characterize the spatial dynamics of malaria [2] and to allocate resources effectively. On the one hand, geostatistical modeling approaches have been used to generate maps of various epidemiological quantities, such as parasite prevalence [3] and intervention impact [4]. These maps derive from methods that interpolate across spatially idiosyncratic data sources, providing a spatially smoothed estimate of epidemiological metrics that are relevant for spatial targeting of interventions. On the other hand, there are important aspects of spatial malaria epidemiology that interpolation methods cannot capture. First, different assumptions about connectivity can lead to differences between the areas that are identified as hotspots for incidence and those that are hotspots for transmission [5], with the latter being the most ideal targets for intervention. Second, thinking beyond all but the most local scales, there are a myriad of ways that control efforts across different areas could be coordinated. By characterizing structure in malaria transmission across different locations, groups of locations can be identified such that transmission within groups is greater than it is between them [6, 7]. Combined with transmission models that take into account numerous nonlinear feedbacks between control and transmission [8, 9] and that are capable of accounting for location-specific intervention packages and their impacts [10, 11], these approaches could in theory suggest an optimal elimination strategy. In practice, there are shortcomings of both currently available data and models.

Quantifying connectivity is one of the most important aspects of characterizing spatial dynamics of malaria, yet it can be one of the most vexing. Call data records and other novel data sources on human travel have offered a great deal of hope in recent years [5, 7, 12], but those methods have their challenges, including variable cell tower densities, mobile phone market fragmentation, and possible disconnects between who is making calls and who is transmitting parasites [13]. Traditional travel survey data may be more directly related to known symptomatic individuals, however these data are often limited in scope and accuracy [14]. Ultimately, models are necessary to appropriately combine information about human mobility with a variety of epidemiological data to arrive at an estimate of how parasite movement is structured. Among other data sources that figure into this process of elucidating patterns of parasite movement, parasite genetic signals may offer some of the richest information about these otherwise elusive patterns of parasite movement. This signal may be easiest to identify in low transmission settings [15] for reasons discussed further below.

Although it was not explicitly spatial, a recent analysis of temporal trends in malaria dynamics highlights the potential value of using transmission models to assimilate parasite genetic and other data to make epidemiologically meaningful inferences [16]. In this work, 24-SNP barcodes from 1,007 parasite samples collected over an eight-year period were used together with a detailed model of transmission dynamics within a city in Senegal to infer that there had been a decline in transmission followed by a rebound. The model and genetic data worked together by determining which transmission scenarios had the highest likelihood of generating patterns consistent with the genetic data. In this example, parasite genetic data offered additional support about temporal trends in support of the epidemiological analysis. We propose that the marriage of parasite genetic data and models in a spatial context may offer unique insights into the epidemiology of malaria.

### Applications of parasite genetics to spatial epidemiology of malaria

Molecular tools may be most valuable when epidemiological information is scarce and/or mobility data is unavailable, to quantify changes in transmission and identify the patterns of pathogen transmission between different locations. Genomic surveillance and phylogenetic analyses that relate the geographic distribution of genetic signals within and between populations have enabled near real-time estimation of transmission chains for non-sexually recombining, rapidly evolving pathogens (e.g. Ebola, influenza) [17, 18]. This nascent field of pathogen phylogeography has provided key insights into the routes of pathogen introductions and spread, particularly for viral diseases. However, directly extending these methods to a pathogen like malaria – a sexually recombining eukaryotic parasite with a complex lifecycle – requires both molecular and analytic advancements that are still at early stages of development. In particular, the malaria parasite *Plasmodium falciparum* undergoes frequent sexual recombination, and is often characterized by multi-genotype infections and low density chronic blood-stage infections that can last for months in asymptomatic individuals. These complexities mean that standard population genetic or phylogenetic approaches do not resolve relationships between parasite lineages effectively [19].

Most national control programs are interested in spatial scales that are operationally relevant: namely, within a country or between countries if they are connected by migration. Population differentiation on international and continental geographic scales can be identified using principal component analysis (PCA), phylogenetic analysis and *F*_*ST*_ [20–24], but these methods are not powered to detect finer-scale differentiation. This is because (1) recombination violates the assumptions underpinning phylogenetic analyses [25]; and (2) PCA and *F*_*ST*_ are influenced by drivers of genetic variation that act on a long time scale (i.e. the coalescent time of parasites) such that, if migration happens multiple times during this time frame, genetic drift can dominate and lead to little or no signal of differentiation among populations [26, 27]. In contrast, methods that exploit the signal left by recombination, rather than treating it as a nuisance factor, may have the power to detect geographic differentiation on spatial scales relevant for control programs. Recombination occurs in the mosquito midgut when genetically distinct gametocytes come together to form a zygote, leading to the production of sporozoites (and hence onward infections) that are highly related. These highly related parasites will tend to have genomes with a high degree of identity over long contiguous blocks, which can be detected given sufficient density of informative markers. Perhaps the simplest measure of genetic similarity is identity by state (IBS), which is defined as the proportion of identical sites between two genomes and is a simple correlate of genetic relatedness between parasites. IBS, however, makes no distinction between sites that are identical by chance and those that are identical due to shared ancestry, making it sensitive to the allele frequency spectrum of the particular population under study. Analyses that are probabilistic and therefore account for identity by chance (e.g. STRUCTURE [28]) provide better resolution, but ultimately linkage disequilibrium-based methods, such as identity by decent (IBD) [29, 30] and chromosome painting [31], which both account for identity by chance and harness the patterns of genetic linkage disequilibrium that are broken down by recombination, are more sensitive to recent migration events and thus can be powerful at smaller geographic scale.

In low transmission settings, such as Senegal and Panama, STRUCTURE as well as IBS (which approximates IBD, albeit with more noise), can often be used to cluster cases together and infer transmission patterns within countries [32–34]. In intermediate transmission settings, such as coastal regions of Kenya and border regions of Thailand, where genetic diversity is higher, IBS, IBD, and relatedness based on chromosome painting have been shown to recover genetic structure over populations of parasites on local spatial scales [27, 35]. However, due to dependence on allele frequency spectra, IBS is not as easily comparable across data sets, and as mentioned above, can be overwhelmed by noise due to identity by chance. Moreover, all of these methods currently have limited support for polyclonal samples. In high transmission settings, the complexity of infection (COI) is very high, making it difficult to calculate genetic relatedness between parasites within polyclonal infections or estimate allele frequencies across polyclonal infections. Methods are available to phase parasite genetic data within polyclonal infections [36], while THE REAL McCOIL [37] has been developed to infer allele frequencies and COI simultaneously, allowing downstream calculation of *F*_*ST*_. However, to fully characterize genetic structure at fine scales in high transmission settings, new methods are needed that estimate IBD and other relatedness measures to infer ancestry between polyclonal infections. Indeed, across all spatiotemporal scales we propose that rather than being defined by the transmission of discrete (clonal) parasite lineages, malaria epidemiology may be best characterized as the transmission of (often polyclonal) infection states.

### Current sequencing strategies for genomic epidemiology of malaria

The use of the population genetic approaches described above will depend on routine collection of parasite genetic data, and must be tailored to the sequencing approach. The discriminatory power of the genotyping method will depend on the local epidemiology and transmission setting. The two most commonly used approaches, relatively small SNP barcodes and panels of microsatellite markers [38], have been extensively used to monitor changes in the diversity and structure of the parasite population. However, signals in these markers may not be sufficient to distinguish geographic origin, and they have limited resolution in certain transmission settings [39, 40]. Increasing the discrimination of each locus will be necessary, and probably more effective than sequencing additional loci, for questions relevant to elimination. Identifying a panel of optimally informative genetic markers to address a specific question remains a major challenge that must balance the cost, throughput, and discriminatory power between (often polyclonal) infections. For example, at fine geographic scales high resolution markers with representative coverage of the genome may be required as compared to studies comparing distant parasite populations; density of sampling infected individuals will also affect the number and type of loci required.

With proper consideration, a parsimonious set of genetic targets may be identified as useful to answer a number of questions with malaria genomics, but these may be context specific, both in terms of the location and the scientific question. Nonetheless, the development of a tool box of genotyping methods tailored to answering questions relevant for transmission on different spatial scales is an important goal. To this end, several ambitious sequencing studies have begun, and over 4,000 *Plasmodium falciparum* genomes sequenced from different transmission settings around the globe (such as the Pf3K Project – https://www.malariagen.net/data/pf3k-pilot-data-release-3) [41–43]. These genetic data are all publicly available, providing a crucial framework to build upon when designing more local, sequence-based epidemiological studies that balance the trade-off between the number of genetic loci evaluated with the quality of the data (e.g. depth of sequence coverage) for each parasite sample. Genomic sequencing methods are evolving rapidly towards high throughput and low-cost deep sequencing approaches that can be done on routinely collected patient samples, allowing for evaluation of even asymptomatic, low density infections e.g. by selective enrichment of parasite DNA [44, 45]. Moving forward, we must ensure that rich metadata are also made easily available in the context of genome sequences, so that links can be made to epidemiological and ecological variables and models.

### Combining data layers to map malaria

In concrete terms, we want to be able to clearly identify if two locations are epidemiologically linked. However, given the current methods available and in development, the complicated lifecycle of the parasite, and epidemiology of malaria, any single data source or method is unlikely to produce a complete picture of the spatial dynamics of malaria parasites. Figure 2 illustrates an analytical pipeline linking different spatially explicit data sets to methods and ultimately interventions, highlighting current uncertainties and the need to take policy-relevant metrics into account when designing sampling frameworks. In particular, we believe that future development should focus on identifying how these different types of data can be combined and integrated to provide a more complete picture of connectivity and transmission dynamics. If we view this problem in terms of a simplified traditional medical statistic, malaria parasite data have a high false negative rate (the analysis mostly underestimates relatedness between parasites) whereas connectivity data inferred from mobile phone data or other proxy measures of travel have a high false positive rate (the analysis mostly overestimates the number of epidemiologically relevant connections). Ideally, additional joint inference methods that combine these data sources would help improve the type I (false positivity rate) and type II (false negativity rate) errors in each type of data.

**Figure 2:**
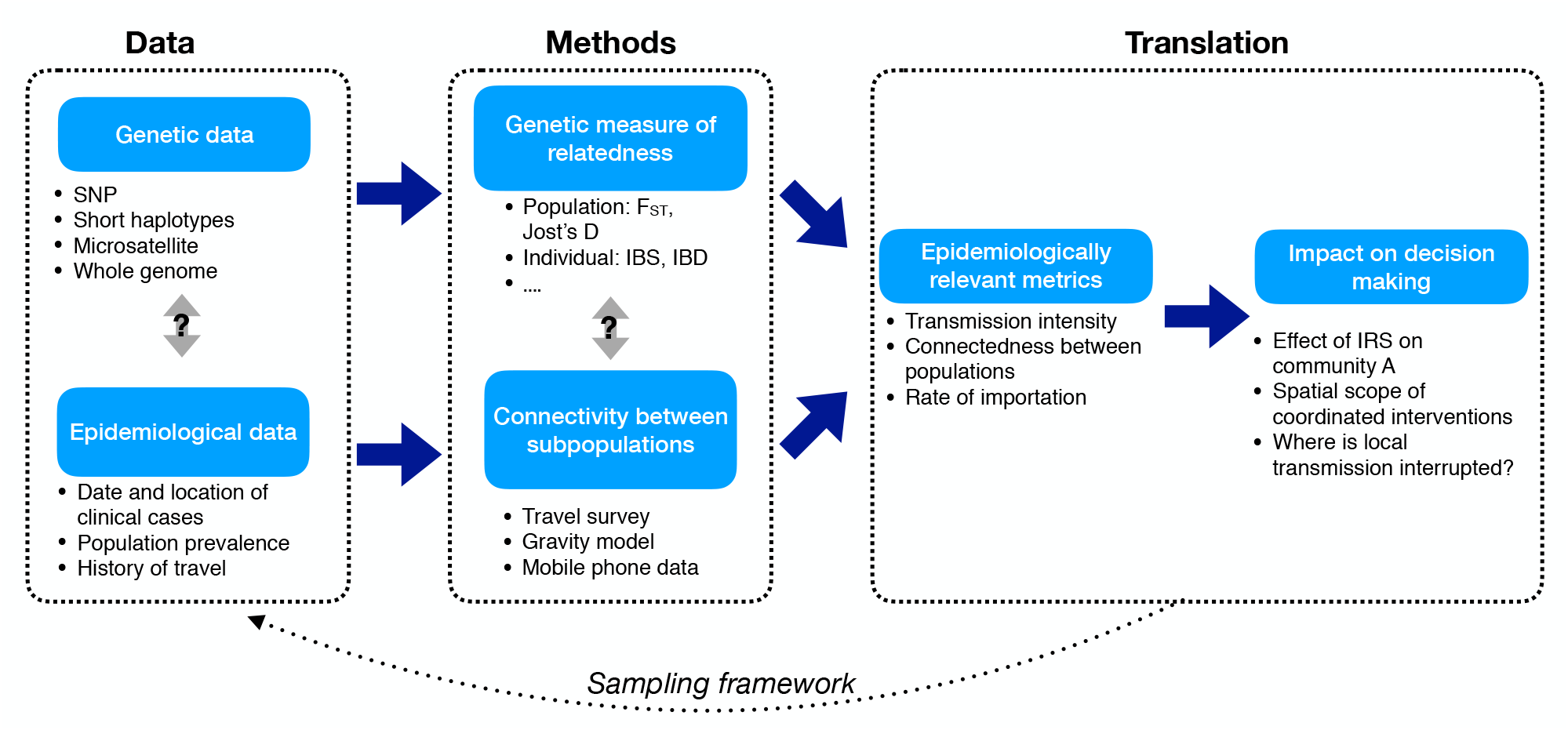
The analysis pipeline. Both genetic and epidemiological data can be collected and analyzed in order to understand parasite flow (with example data sets and methods listed above). Identifying how these two methods can be combined, directly related to policy relevant questions, and translated control measures will require the development of novel inference frameworks and designed studies (sampling framework) across a range of transmission settings.

These new data streams therefore offer great potential, but understanding how to effectively combine them in ways that take into account the biases and strengths of each data type will require significant research investment. Furthermore, making these methods relevant for implementation is a consideration that must be at the forefront of research efforts. For example, the ongoing availability of each data stream, the feasibility of implementing the analytical pipeline in the context of the national control program, and the capacity building required to do so, will ultimately determine the impact of these approaches. This means that tools must provide clear estimates of uncertainty, and will need to be straightforward to use in different contexts, easy to communicate, and generalizable.

## Acknowledgements

This work is supported by Maximizing Investigators’ Research Award for Early Stage Investigators, R35GM124715 (COB, AW, ART), a Wellcome Trust Sustaining Health Grant, 106866/Z/15/Z (COB, AW, ART) (https://wellcome.ac.uk/), the Models of Infectious Disease Agent Study program, cooperative agreement U54GM088558 (COB) (https://www.nigms.nih.gov/Research/specificareas/MIDAS/Pages/default.aspx),<u>and</u> the Bill and Melinda Gates Foundation OPP 1132226 (TAP, BG, ST), OPP 1110495 (TAP). BG is a Chan Zuckerberg Biohub investigator. AW is supported by a Career Award at the Scientific Interface from the Burroughs Wellcome Fund. RV is funded by a Skills Development Fellowship: this award is jointly funded by the UK Medical Research Council (MRC) and the UK Department for International Development (DFID) under the MRC/DFID Concordat agreement and is also part of the EDCTP2 programme supported by the European Union

## Competing Interests

None of the authors report any competing interests.

